# Abundance of human T-cell epitopes in microbial proteomes

**DOI:** 10.1101/2023.11.29.568792

**Authors:** Vasilisa A. Kovaleva, Ivan V. Zvyagin

## Abstract

Molecular mimicry, the structural similarity between self and foreign antigens, is considered as a key factor in post-infectious autoimmunity. While certain examples of molecular mimicry are well studied, a comprehensive analysis of its prevalence and impact on the type of T cell response to such self is absent. In this work, we comprehensively studied the frequency of molecular mimicry between human and microbial T-cell epitopes. We performed an *in silico* analysis of the occurrence of T-cell epitopes originating from different sets of proteins: normal self epitopes, proteins involved in autoimmunity, and cancer neoantigens, in the proteomes of commensal and pathogenic microbiota. We show a significant overlap between repertoires of human T-cell epitopes and predicted epitopes from proteins of both commensal and pathogenic microbiota: the counterparts for over 90% of human HLA-I and 5% of HLA-II ligands were found in the microbial proteomes. HLA-II epitopes derived from the proteins involved in autoimmunity were more frequent in microbiota compared to normal self, suggesting a potential deleterious effect of molecular mimicry, while we did not observe this effect for HLA-I epitopes. Cancer epitopes were less frequent in microbiota compared to normal self of epitopes, implicating potential cancer escape from cross-reactive T cells specific to microbial antigens. Together, our results show that molecular mimicry might have a general pro-inflammatory effect on similar self epitopes, though much in this field remains to be explored.

## Introduction

Tolerance to self is crucial for proper immune functioning without autoimmunity. During negative selection in the thymus, developing naive T cells with TCRs recognizing epitopes from self-antigens, are eliminated via apoptosis or differentiate into regulatory T cells (Treg), which can suppress the immune response. Recent works showed that during development in thymus, each immature T cell can interact with ∼10^3^–10^5^ self-epitopes, which is at least 10 times lower than the total potential pool of self-peptides (Wortel et al., 2020), suggesting that naive T cells with potentially autoreactive TCRs can pass through the negative selection (Gallegos and Bevan, 2006). The limitations of central tolerance are complemented by peripheral tolerance involving suppression of T-cell response by regulatory T cells, induction of anergy state by cytokines and/or in absence of necessary co-stimulus or by action of inhibitory checkpoint molecules, to prevent autoimmunity by T cells which escaped elimination (ElTanbouly and Noelle, 2021). Both, the central (thymic selection) and peripheral tolerance can also be directed towards foreign antigens derived from food (Faria and Weiner, 2005) or commensal microbiota (Nutsch and Hsieh, 2012).

Commensal microbiota residing in our body are represented by ten times more number of cells than our organism. The estimated number of unique proteins expressed by the microbiome also exceeds the human proteome by an order of magnitude (Ley et al., 2006). Our immune system is thus constantly exposed to a massive amount of antigens derived from microbiota which are mostly different from self and should be recognised as foreign with appropriate implications. Central and peripheral tolerance work together to prevent or balance the inflammatory response to the microbiome allowing immune control. As it was recently shown, peptides from commensal microorganisms are transported by peripheral dendritic cells to the thymus to be presented during the selection of T cells (Klein et al., 2014; Zegarra-Ruiz et al., 2021).

Microorganisms can promote suppressive conditions of the immune system. Naive T cells in the colon interacting with commensal microbiota can differentiate into Tregs rather than effector T cells (Lathrop et al., 2011). Some bacterial strains can produce tolerogenic metabolites such as short-chain fatty acids (Arpaia et al., 2013) and polysaccharide A (Round and Mazmanian, 2010), and their properties are now actively explored for autoimmunity treatment (Golpour et al., 2023). These observations led to the “extended self” concept. Today many researchers view the microbiota this way (Berer and Krishnamoorthy, 2012; Rice and Palm, 2016). Thus while dissimilarity to the self-epitopes might be used as a predictor for peptide immunogenicity due to negative selection process (Richman et al., 2019), the immune tolerance is not strictly restricted to self-antigens, shows some flexibility, and extends to some foreign antigens, including our commensal microbiome (Zhang et al., 2020).

Structural similarity of antigens can lead to similar response by the immune system to those antigens. Microbial antigens from certain infectious agents exhibit a close resemblance to self-antigens. During inflammation, this similarity potentially can disrupt peripheral tolerance, leading to an autoimmune reaction against the self-antigen (Gugliesi et al., 2021; Lucchese and Flöel, 2020a, 2020b; Luo et al., 2018; Tengvall et al., 2019). Structural similarity between foreign and self-antigens is termed molecular mimicry and is one of the leading mechanisms of post-infectious autoimmunity development (Rojas et al., 2018). Similarly, if some microbial antigens resembled tumor antigens, conceivably, the immune response against the pathogen would trigger an antitumor response. It was demonstrated that T cell cross-reactivity between microbiome- and cancer-derived antigens facilitates anti-tumor immunity (Bessell et al., 2020; Fluckiger et al., 2020). However, it is still unknown how common molecular mimicry is, either between microbial and self-peptides or between microbial and tumor neoantigens.

In this work, we studied the frequency of molecular mimicry between human and microbial T-cell epitopes. We investigate the occurrence of T-cell epitopes originating from different sets of proteins: normal self, proteins involved in autoimmunity, and cancer neoantigens, in the proteomes of the commensal and pathogenic microbiota.

We show that molecular mimicry of human HLA ligands with microbial epitopes is fairly common: matches for approximately 90% of human HLA-I ligands and for 5% of human HLA-II ligands can be detected in microbial proteomes, showing that such similarity itself is not enough to trigger autoimmunity. HLA-II epitopes derived from the proteins involved in autoimmunity are more frequent and cancer neoantigens are less frequent in microbiota compared to the control group of normal self-epitopes. The last finding implicates potential cancer escape from cross-reactivity with T cells specific to microbiota. Together, the results suggest little if present tolerogenic effect of similarity to commensals when considering the whole range of self T-cell epitopes.

## Materials and Methods

### Epitopes

In our study, we used four epitope types: (1) HLA ligands obtained from HLA Ligand Atlas (Marcu et al., 2021) (further referred to as “HLA ligands”); *in silico* predicted epitopes from (2) randomly selected human proteins (“self epitopes”) and (3) autoimmune proteins (“autoimmune epitopes”), and (4) epitopes predicted from cancer neoantigens (“cancer epitopes”). HLA-I epitopes were 9 amino acids long, and HLA-II epitopes 15 amino acids long, since these were the most abundant epitope lengths for their types in HLA Ligand Atlas.

All selected 9- and 15-mers were predicted for binding to HLA-I or HLA-II using NetMHCpan-4.0 and NetMHCIIpan-4.0 respectively (Reynisson et al., 2020). HLA-I epitopes were predicted for binding to HLA-I supertype representative alleles: HLA-A01:01, HLA-A02:01, HLA-A03:01, HLA-A24:02, HLA-A26:01, HLA-B07:02, HLA-B08:01, HLA-B27:05, HLA-B39:01, HLA-B40:01, HLA-B58:01, HLA-B15:01. For HLA-II, we used the most frequent human HLA-II alleles according to (Jawdat et al., 2020): DRB1_0701, DRB1_0301, DRB1_1302, DRB1_1501, DRB1_1301, DRB1_0403, DRB1_0405. We used default thresholds for binding affinity, only strong binders were selected for further analysis.

#### HLA ligands

To obtain a set of HLA ligands, we used data from the HLA Ligand Atlas database (Marcu et al., 2021). 34,000 and 17,000 ligands for HLA-I and HLA-II, respectively, were randomly selected from sets for different tissues. Then, we predicted their binding with HLA. We selected 25,984 HLA-I and 5,733 HLA-II epitopes as strong binders. A negligible amount of resulting HLA-II epitopes contained HLA-I epitopes within their sequences: 251 HLA-I epitopes were substrings of HLA-II epitopes, and 338 HLA-II epitopes contained at least one HLA-I ligand.

#### Autoimmune epitopes

To generate a set of autoimmune epitopes, we retrieved 2,612 autoimmunity-related epitopes from the Immune Epitopes Database or IEDB (Vita et al., 2019), using the following search parameters: “epitope” - linear, “epitope source” - *Homo sapiens*, “assay” - T-cell assay, “disease” - autoimmune. We recovered 1,012 reference amino acid sequences of proteins containing these epitopes (UniProt database, release-2021_04) (The UniProt Consortium, 2021), and *in silico* generated all possible 9- and 15-mers using the sliding window approach. After HLA-binding prediction, we selected 9,304 HLA-I and 4,746 HLA-II strong binders representing 557 and 503 proteins respectively (**Supplementary Material 1**).

#### Self epitopes

To generate self epitopes, we retrieved a list of 15,262 human proteins containing HLA ligands within their sequence confirmed by HLA Ligand Atlas. From this list, we excluded 1,275 proteins that were present in our list of proteins used to generate autoimmune epitopes and 288 proteins associated with autoimmunity according to DisGeNET (Piñero et al., 2020) (see **Supplementary Material 2**). From the remaining list, we randomly chose three sets, each containing 600 unique protein sequences. From each sequence, all possible 9- and 15-mers were *in silico* generated. After HLA-binding prediction, sets of 25,246, 29,352, and 26,521 HLA-I epitopes, and 54,634, 58,690, and 55,819 HLA-II epitopes were selected as strong binders.

#### Cancer epitopes

We have randomly chosen three sets of cancer neoantigen epitopes from The Cancer Genome Atlas (https://www.cancer.gov/tcga). For each cancer epitope, we identified the source protein and recovered an epitope sequence without mutation based on the reference sequence from UniProt. We will refer to them as “cancer original” epitopes. Subsequently, we randomly mutated *in silico* the cancer original epitope sequences based on a human mutation profile (Vitkup et al., 2003). The set of such mutated epitopes is referred to as “cancer mutated” here. For each epitope, we predicted binding with the list of HLA-I alleles. This way, we generated three groups of cancer epitopes: (1) “cancer” epitopes, *i.e.* real cancer neoantigen epitopes from TCGA, (2) “cancer original” epitopes *i.e.* reference sequences without cancer mutations, and (3) “cancer mutated” epitopes, *i.e.* randomly mutated original epitopes. All epitopes whose counterparts were predicted as weak- or non-binders were excluded. Consequently, we obtained three sets of 4,929, 4,920, and 4,944 cancer epitopes.

To better estimate the variance in similarity with microorganisms, for self epitopes and cancer neoantigen epitopes, we used three sets of epitopes for each group. To ensure the diversity of the sets within one epitope group, we filtered out those having matches with Hamming distance ≤2 for HLA-I or ≤4 for HLA-II between different sets (*e.g.* “self-1”, “self-2” and “self-3”) of one group (“self” or “cancer”).

### Proteomes

To study the similarity between human and microbiota potential T-cell epitopes, we analyzed proteomes of three different groups of microorganisms: (1) Commensal microorganisms (commensal bacteria), (2) Pathogenic microorganisms (pathogenic bacteria, fungi, and viruses), and (3) Neutral microorganisms (soil bacteria). Reference proteomes of the bacteria were obtained from the UniProt database with the “Complete Proteome Detector” parameter equal to “Standart.” All included genera are listed in **Supplementary Material 3**.

#### Commensals

188 bacteria genera of commensals were selected from the GMRepo database (Dai et al., 2021) based on their prevalence: only genera found in more than 5% of the samples with abundance higher than median abundance among all samples were included.

*Paraprevotella*, *Prevotella*, *Bacteroides*, *Dialister*, *Megasphaera*, *Eubacterium*, *Alkaliphilus*, *Clostridium*, *Ruminiclostridium*, *Faecalibacterium*, *Coprococcus*, *Butyrivibrio*, *Anaerostipes*, *Pseudobutyrivibrio*, *Roseburia*, *Dorea*, *Blautia*, *Lachnoclostridium* genera, and their close relatives (from the closest monophyletic groups (**Supplementary Figure 1**) were selected as short-chain fatty acid (SCFA) producing bacteria based on literature (Deleu et al., 2021).

#### Pathogens

The list of 38 genera of pathogenic bacteria was obtained from the PATRIC database (Davis et al., 2020). The list of 28 genera of pathogenic fungi was obtained from the MSD Manual (Vergidis, 2023). As a source of viral proteomes, we used the ViPR database (Pickett et al., 2012) with the following search criteria: “host” – “human”, “data to return’’ – “protein”, and “complete sequences only”. The metaproteome of the whole viral family was used in each case: in total 17 families’ metaproteomes were included.

#### Neutrals

170 soil bacteria genera were randomly selected from (Fulthorpe et al., 2008), Proteomes included in the commensal bacteria group were not selected as neutrals.

### Workflow

The pipeline (**Figure 1**) for human epitope search in a microbial proteome consisted of the following steps:

1. Generation of k-mer database from the proteome;
2. Search for the epitope match in the k-mer database within a Hamming distance of 2 (for HLA-I epitopes) or 4 (for HLA-II).
3. HLA binding prediction for the sequence (“match”) matching with the epitope;
4. Filtration of resulting matches based on binding prediction. Matches not binding to at least one HLA allele present in the HLA allele list for the original epitope were filtered out.

**Figure 1.**
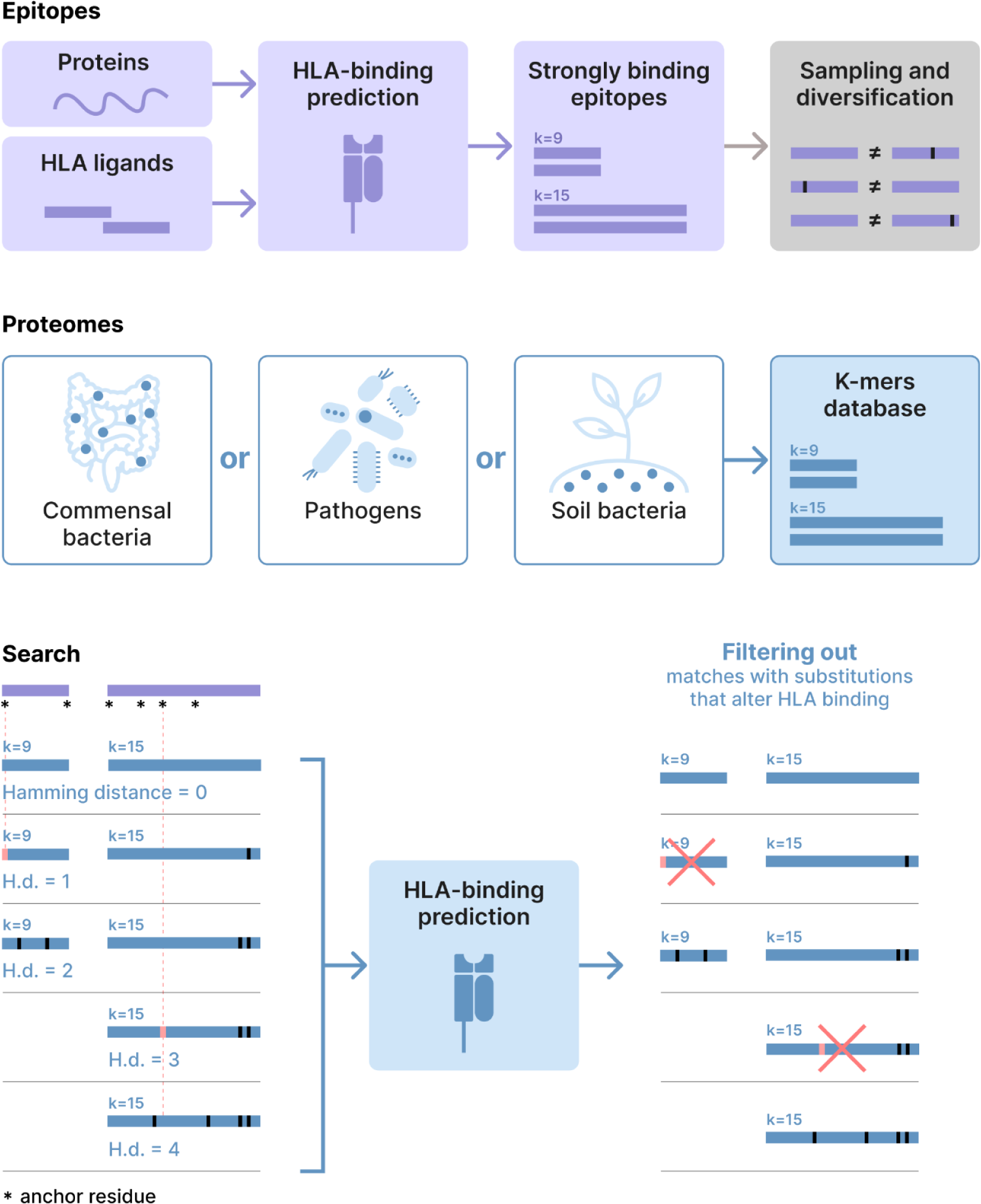
Workflow for search of human T-cell epitopes in microbial proteomes. Peptides of length 9 for HLA-I epitopes and 15 for HLA-II epitopes were retrieved from the HLA-ligandome database or human protein sequences and predicted for HLA binding. Only strong binders were selected for further analysis. A diversification procedure was subsequently applied to self epitopes and cancer neoantigen epitopes (see the Materials and Methods section for details). From the proteomes of microorganisms the database of all possible 9- and 15-mers was generated. Then, we searched for the epitopes in the database allowing up to 2 mismatches for HLA-I epitopes and up to 4 mismatches for HLA-II epitopes. The matching 9- or 15-mers from microorganism’s proteomes were further predicted for HLA-binding to get rid of the matches with substitutions altering HLA binding. The black line in the match (blue) indicates a substitution in comparison to an original epitope (violet). The pink line indicates a substitution altering the binding to HLA.

The code for the k-mer database generation and search steps is available at vasilisa-kovaliova/MolecularMimicry.

We used three measures of similarity between human and microbiota potential T-cell epitopes:

1. Findability is the ratio of epitopes with matches to the total number of epitopes in the set (%). For self epitopes, we also generated a distribution of findability values for 1000 random subsamples. The size of the random sample corresponded to the size of the set of the epitope group of interest (autoimmune or cancer epitopes sample).
2. Number of matches per epitope represents the number of times one epitope was detected in all proteomes of the particular microorganism’s group. The number of matches per epitope reflects the “epitope occurrence” in the microbiota metaproteome.
3. Number of matches per genus is the number of epitopes found in the proteome of one bacterium. Since the number of searched epitopes influences this measure, it was normalized by the number of searched epitopes of the same group. This measure was applied only to the search in Commensals and Neutrals since Pathogens included not only pathogenic bacteria but also fungi and viruses.

## Results

### HLA ligands have high similarity to potential microbial epitopes

To study the similarity at the T-cell epitope level between human and microbiota, we searched for HLA-I and HLA-II ligands obtained from the HLA Ligand Atlas in proteomes of Commensals, Pathogens, and Neutrals (see Materials and Methods section for details). To account for polymorphisms in human epitopes and the potential TCR ability to recognize similar but not identical epitopes, we allowed up to 2 substitutions in matches to HLA-I epitopes and up to 4 substitutions for HLA-II epitopes. We only included epitopes of the most prevalent lengths in HLA Ligand Atlas: 9 amino acids for HLA-I epitopes and 15 amino acids for HLA-II epitopes.

To evaluate similarity, we used two metrics: findability, calculated as the ratio of the number of epitopes having matches in microorganisms to the total number of epitopes, and the number of matches per epitope, calculated as the number of times an epitope was found in a metaproteome.

Results for HLA-I (n=25,984 ligands) and HLA-II (n=5,733 ligands) are displayed in red and blue respectively.

For HLA-I epitopes, we observed a surprisingly high level of similarity with all three tested microorganism groups. Specifically, over 95% of HLA-I epitopes were found at least once in Сommensals, 87% in Pathogens, and more than 95% in Neutrals with up to 2 substitutions allowed per 9 positions (**Figure 2A**). However, the findability of exact matches was not so high: only 0.9% of HLA-I epitopes had exact matches in the Commensals, 1.6% in Pathogens, and 0.78% in Neutrals.

**Figure 2.**
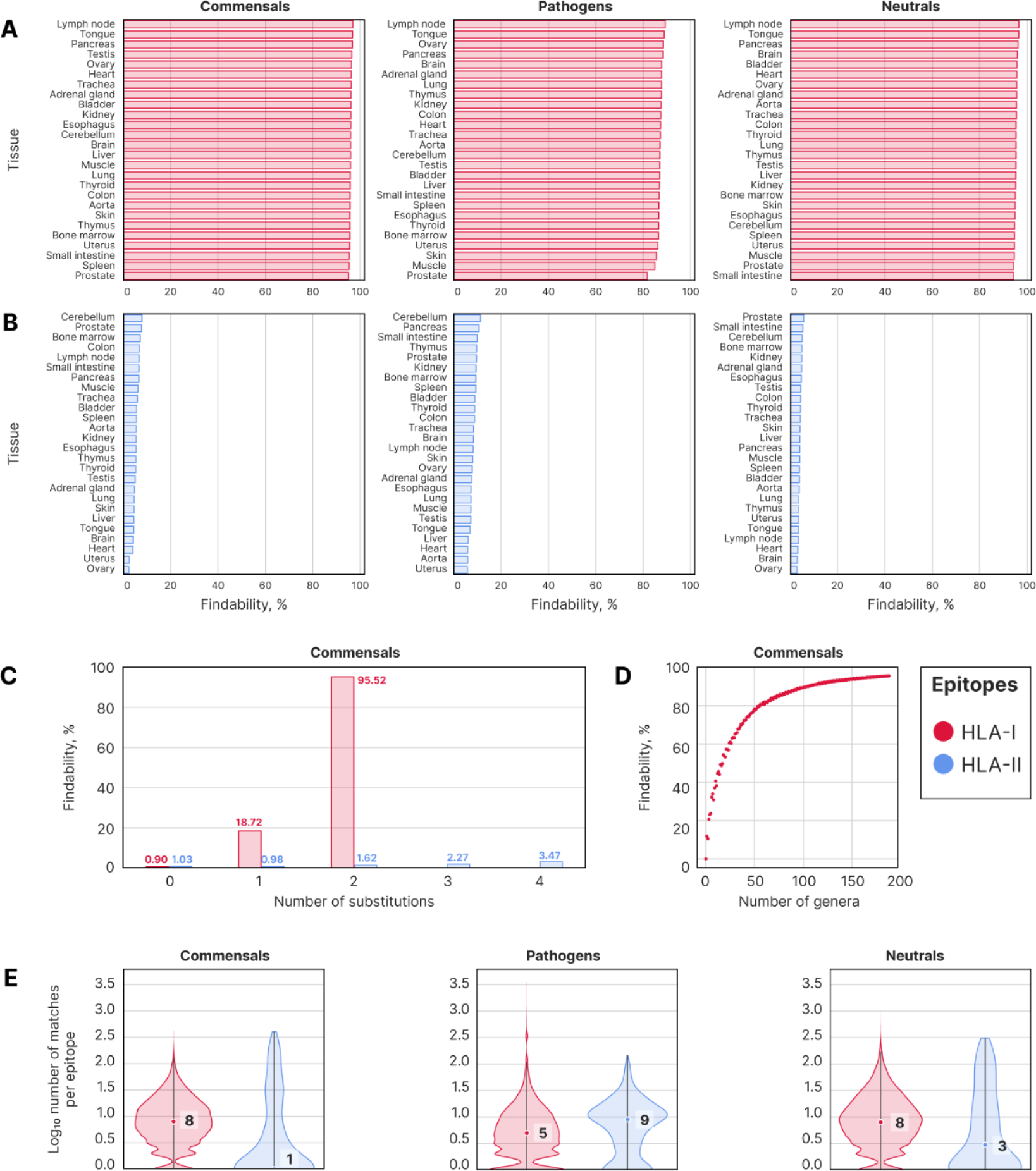
Occurrence of HLA-I and HLA-II ligands in the proteomes of Commensals, Pathogens, and Neutrals. Findability of HLA-I ligands (**A**) and HLA-II ligands (**B**) derived from different tissues. **C**: Distribution of findability depending on the number of allowed substitutions. Note, that findability values for one epitope type do not sum up to 100%, as a match for an epitope could be found with both 0 and 1 substitutions simultaneously. **D**: Findability distribution depending on the number of searched proteomes. For each possible number of bacterial genera, findability was measured 20 times with randomly selected genera. **E**: Number of matches per epitope. The median number of matches per epitope is shown for each violin.

In contrast, HLA-II epitopes showed much less similarity to microbiota: 5% of HLA-II epitopes were detected in Commensals, 9% in Pathogens, and 4% in Neutrals (**Figure 2B**). Such a drastic difference in findability between HLA-I and HLA-II epitopes can be attributed to the difference in their length. Again, the findability of exact matches was considerably lower and did not change strongly: 1% of HLA-II epitopes had exact matches in Commensals, 1.65% in Pathogens, and 0.34% in Neutrals (**Figure 2C**). Only a few HLA II epitopes contained sequences of HLA-I epitopes, so this low overlap had almost no influence on the findability of epitopes (**Supplementary Figure 2B**).

Epitopes across all tissues had comparable findability (**Figures 2A and 2B**), showing the absence of contribution of particular tissues to the results due to tissue-specific expression. Tissues more exposed to foreign antigens (e.g., barrier tissues and the digestive system) showed the same similarity level as other tissues, suggesting the minimal influence of potential contamination of eluted HLA-ligands by bacterial peptides on the measured similarity.

Since some genera could contribute to the findability more than others, we analyzed how findability depended on the number of bacterial proteomes. Findability grew logistically with the number of randomly selected bacteria genera (**Figure 2D**), and 50 genera were enough to reach a findability of 80% for HLA-I epitopes.

On average, an HLA-I epitope was detected 8 times in Commensals, 5 times in Pathogens, and 8 times in Neutrals. For HLA-II epitopes, the median number of matches per epitope was lower: 1 in Commensals, 9 in Pathogens, and 3 in Neutrals. Notably, HLA-I epitopes were less frequently identified in Pathogens in contrast to HLA-II epitopes, which were more frequently detected in Pathogens compared to other microorganism groups (**Figure 2E**).

For certain epitopes, the number of matches in microbiota was exceptionally high. Some HLA-I epitopes were detected over 2,400 times in Commensals, more than 3,600 in Pathogens, and more than 2,000 times in Neutrals. For HLA-II epitopes, the maximum was lower, with the highest number of matches per epitope reaching 400 in Commensals, 140 in Pathogens, and 300 in Neutrals (**Supplementary Figure 2C**). The number of matches per epitope was linearly dependent on the number of bacteria genera (**Supplementary Figure 2A**).

Overall, we found unexpectedly high similarity between human HLA ligands and microbiota. The similarity at the T-cell epitope level to different microbiota groups, though, potentially could have different consequences for the recognition of human epitopes by the immune system. Generally, the immune system is tolerant towards commensal bacteria, therefore, the similarity of human epitopes to epitopes derived from commensals can have a tolerogenic effect (**Figure 3A, left panel**). On the contrary, pathogens usually lead to an inflammatory immune response, so similarity to their epitopes might have an inflammatory effect (**Figure 3A, right panel**). In further analysis, we evaluated the consequences of similarity to commensals and to pathogens using three epitope groups: autoimmune epitopes, neoantigens, and self epitopes.

**Figure 3.**
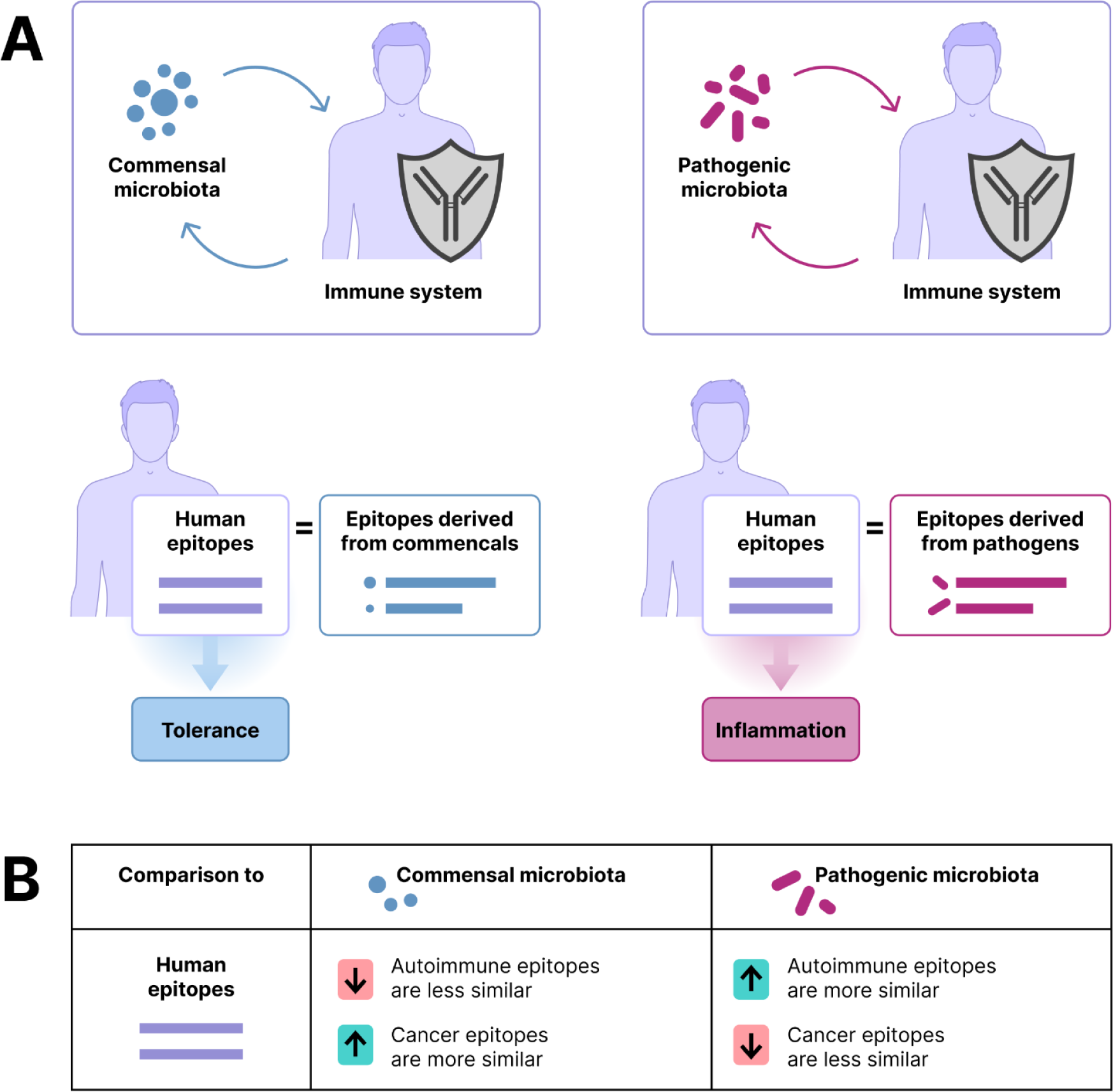
Hypothesis: potential consequences of sharing of T-cell epitopes between human and microbiota. **A:** Commensal microbiota is tolerated by the immune system, while contact with pathogenic microbiota leads to an inflammatory immune response. **B:** Based on this, we hypothesized that human epitopes similar to commensal-derived epitopes are likely to be tolerated by the immune system, while those similar to pathogenic epitopes may be involved in inflammation. To test this hypothesis, we compared the similarity to commensals and pathogens between autoimmune, cancer, and normal self-epitopes. We suggested that a breach of tolerance towards self-epitopes implicated in autoimmunity may be associated with greater similarity to pro-inflammatory microbiota and/or lower similarity with commensals. In contrast, cancer epitopes were expected to show less similarity to pathogens and greater similarity to commensals due to evolution of cancer to escape the cross-reactivity with inflammation-driving microbiota.

We hypothesized that 1) epitopes involved in autoimmunity would be more abundant in proteomes of pathogenic microbiota and/or lower in proteomes of commensals, and 2) cancer neoantigens would be less frequent in proteomes of pathogens and/or more frequent in commensals **(Figure 3B)**. Self epitopes were included as a control group. We measured the similarity between these three epitope groups and three microbiota groups: Commensals (commensal bacteria), Pathogens (pathogenic bacteria, viruses, and fungi), and Neutrals (soil bacteria).

### HLA-I and HLA-II epitopes involved in autoimmunity have different frequencies in microbiota relative to the control self

We compared the similarity of autoimmune epitopes with the similarity of self epitopes to different microbiota groups. Autoimmune epitopes were generated *in silico* from proteins containing confirmed epitopes involved in autoimmunity, as listed in the Immune Epitope Data Base (IEDB, Vita et al., 2019). Only epitopes predicted as strong binders for a set of the most common HLA alleles were used.

To maintain consistency and minimize potential biases between epitope groups, self epitopes were also generated *in silico*, instead of using HLA ligands from real immunopeptidome data. They were derived from a random set of human proteins containing confirmed HLA ligands according to the HLA Ligand Atlas. Proteins involved in autoimmunity according to the IEDB search were excluded from the set. As we expected, *in silico* generation of epitopes introduced bias to the results, as both findability and the number of matches per epitope were slightly lower for *in silico* predicted epitopes compared to HLA ligands (**Figure 1A, B, Supplementary Figure 2A, B**). To reliably estimate the baseline similarity between self epitopes and microbiota, we used three diversified *in silico* predicted self epitope sets instead of one. The similarity level to the same microbiota group measured via findability and number of matches was notably consistent across all three sets (**Figure 4A, 4B, Supplementary Figure 2A, B**).

**Figure 4.**
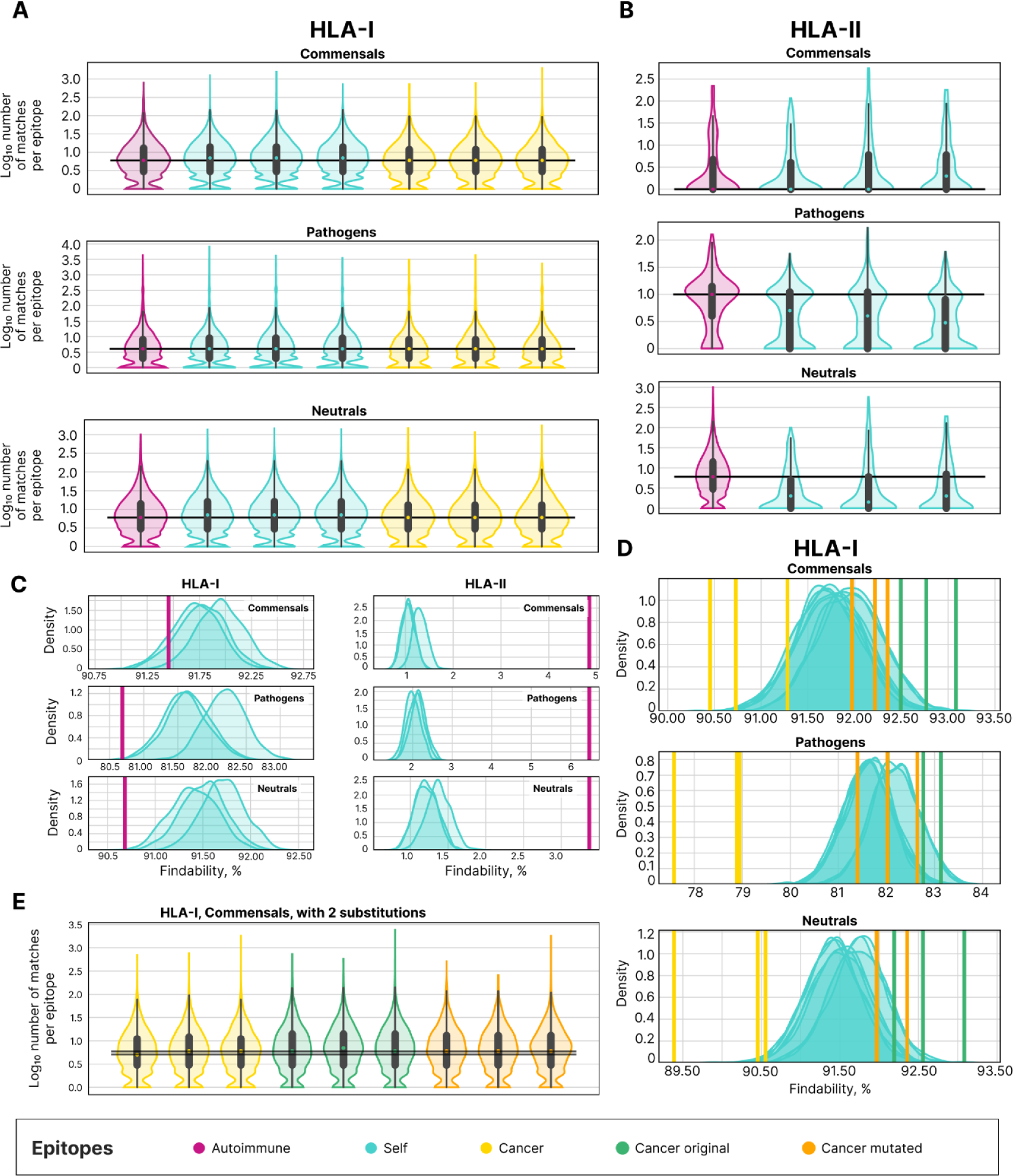
Occurrence of self, autoimmune, and cancer epitopes in the proteomes of Commensals, Pathogens, and Neutrals. Number of matches per HLA-I (**A**) or HLA-II (**B**) epitope. The gray line shows the position of the median number of matches per autoimmune epitope. **C**: Distribution of findability values for 1000 random samples of self epitopes compared to the findability of autoimmune epitopes (**C**, shown by purple vertical lines) or different cancer epitopes (**D**, shown by yellow, orange, and green vertical lines). For exact values of findability see **Supplementary Table 1**. **E**: Number of matches per epitope for cancer, cancer original, and cancer mutated epitopes, with the number of substitutions equal to 2. Two horizontal gray lines show the position of the maximal and minimal median number of matches per epitope for three cancer epitope sets. The sample sizes are as follows: 25,246, 29,352, and 26,521 HLA-I self epitopes, 54,634, 58,690, and 55,819 HLA-II self epitopes; 9,304 HLA-I and 4,746 HLA-II epitopes in the autoimmune sets; 4,929, 4,920, and 4,944 HLA-I epitopes in cancer sets.

Findability for HLA-I autoimmune epitopes was lower than that for self epitopes across all three microbiota groups (**Supplementary Figure 3A**). Since the set of autoimmune epitopes was smaller than each of the sets of self epitopes (9,304 autoimmune HLA-I epitopes and 25,246, 29,352, and 26,521 HLA-I epitopes the three sets of self epitopes), we subsampled self epitope sets to the size of autoimmune epitope set 1000 times, generating a distribution of findability values (**Figure 4C**). In Commensal microbiota, the findability for HLA-I autoimmune epitopes was higher than for only 8.3% of self subsamples (averaged across three self sets). In Pathogens, it was higher than 0.1% and in Neutrals 0.07% of self subsamples (**Supplementary Table 1**).

The number of matches per HLA-I autoimmune epitope was also generally lower than for self epitopes across all microbiota groups (**Figure 4A**). The median number of matches per autoimmune epitope was equal to 6 in Commensals, 4 in Pathogens, and 6 in Neutrals, in comparison to 7 in Commensals, 4 in Pathogens, and 7 in Neutrals for self epitopes. The highest number of matches per epitope was nearly 700 in Commensals, 4,200 in Pathogens, and 1,000 in Neutrals, which is generally lower than for self epitopes (700-1,200 in Commensals, 3,000-8,000 in Pathogens, 1,300-1,400 in Neutrals across three self sets).

In contrast, we got opposite results for HLA-II autoimmune epitopes. The findability for them was considerably higher than for self epitopes (**Supplementary Figure 3B**). As the number of HLA-II self epitopes exceeded that of HLA-II autoimmune epitopes (4,746 autoimmune HLA-II epitopes and 54,634, 58,690, and 55,819 HLA-II epitopes in the sets of self epitopes), again we generated a distribution of findability values for self epitopes (**Figure 4C).** The findability of the autoimmune epitopes was significantly higher compared to 100% of random samples (p-value < 0.001).

Similarly, the number of matches per HLA-II autoimmune epitope was generally higher than for self epitopes (**Figure 4B**). In Commensals, this difference was not observed, with a median number of matches per epitope of 1 for both autoimmune and self epitope sets. However, in Pathogens and Neutrals, the difference was more pronounced: median numbers of matches per epitope were 10 vs 5 for autoimmune and self epitopes in Pathogens, and 3 vs 2 for autoimmune and self epitopes in Neutrals (for self epitopes a maximal median number of matches per epitope from three sets was taken into comparison).

In summary, the results for HLA-I and HLA-II epitopes demonstrated opposite but consistent trends across all three microbiota groups. HLA-I autoimmune epitopes were less similar to microbiota than self epitopes, whereas HLA-II autoimmune epitopes were more similar.

### Cancer epitopes exhibit lower similarity with microbiota compared to self epitopes, cancer original, and cancer mutated epitopes

Cancer neoantigens are another group of self-derived antigens for which a different pattern in similarity with microbiota might be expected due to the nature of the immune response. We hypothesized that cancer might evolve to avoid similarity with pathogenic microbiota if this similarity leads to elevated chances of involvement in inflammation and/or to gain similarity to commensal microbiota if it provides tolerance from the immune system. To test this idea, we compared the similarity of the cancer epitope set (predicted from randomly selected neoantigens, see Materials and Methods for details) to the proteomes of different microbiota groups with the similarity of the self epitopes.

The differences in the similarity between self and cancer epitopes can be explained in three ways: (1) cancer epitopes can originate from proteins that have a different similarity level to microbial proteins compared to baseline; (2) cancer mutations affect the similarity in a non-directional manner; (3) similarity to microbiota guide cancer evolution on T-cell epitope level. To distinguish between these scenarios, for each cancer epitope its original human epitope without cancer mutation was recovered and then *in silico* mutated using a human mutational profile (Vitkup et al., 2003). Hence, each cancer epitope had two counterparts: cancer original epitope (represented “self”) and cancer mutated epitope (represented randomly mutated “self”). To control for variance due to sampling of neoantigens and self epitopes, we used three diversified cancer and self epitope sets (see Materials and Methods for details) as in the previous section.

Cancer epitopes showed a decreased findability compared to self and autoimmune epitopes (**Supplementary Figure 3A**). Since each set of cancer epitopes had fewer epitopes than each set of self epitopes (∼4,900 and ∼25,000 HLA-I epitopes in each set of cancer and self epitopes, respectively), we obtained a distribution of findability values for subsets of self epitopes equal to the size of cancer epitope set (**Figure 4D**). Cancer epitopes showed lower findability in >=97,8% cases for Commensals, Pathogens, and Neutrals, respectively (**Figure 4D**, yellow; **Supplementary Table 1**). In contrast, original cancer epitopes had higher findability than 98.9%, 86.2%, and 94.9% of self epitopes subsets in Commensals, Pathogens, and Neutrals, respectively (**Figure 4D**, green). Random mutation of cancer original epitopes did lower the findability (81.5, 60%, and 94.34%), but it still did not fall to the findability of cancer epitopes (**Figure 4D**, orange), indicating that random mutations can not be solely responsible for this effect.

Number of matches per epitope was lower for cancer epitopes compared to the baseline self in all three groups of proteomes (**Figure 4A**). The median number of matches for cancer epitopes was 5 in Commensals, 2 in Pathogens, and 5 in Neutrals, which was smaller than the observed values for all three self sets in Commensals, Pathogens, and Neutrals (6, 3, and 6 respectively).

Given the method we used to generate cancer original and mutated epitopes from cancer epitopes, these epitopes frequently relate to each other with Hamming distance of 2. Thus this somewhat increases the difference in the number of matches between cancer original set and the two others. To get rid of it, we additionally considered only those matches having exactly 2 substitutions. After this procedure, the findability of cancer epitopes was still lower than the findability of cancer original and cancer mutated epitopes (**Supplementary Figure 3C**). The median number of matches per cancer epitope (5 and 2) was lower than that of cancer original (6 and 3) and mutated epitopes (6 and 3) in Pathogens and Neutrals, respectively (**Supplementary Figure 3D**), but there was no difference in Commensals (the median number of matches per epitope was equal to 6 for cancer, cancer original, and cancer mutated epitope sets) (**Figure 4E**).

To summarize, cancer epitopes were consistently less similar to microbiota than normal self epitopes independently of the microbiota group. Seemingly, this can not be attributed solely to mutations because randomly mutated self epitopes, recovered from original cancer epitopes, did not show such a significant drop in similarity.

### SCFA-producing bacteria do not differ from other commensals in the frequency of autoimmune and cancer epitopes

The number of matches per epitope reflects the occurrence of the epitope in the microbial metaproteome. It can be suggested that commensal microbiota can be tolerated by our immune system better than free-living microorganisms. Also, some bacteria can actively induce tolerance to their antigens due to production of specific metabolites. To detect this potential tolerogenic effect at T-cell epitope level, we estimated the number of matches per proteome of a bacterial genus from the Commensals and Neutrals groups. Unlike findability and the number of matches per epitope, this measure is influenced by the size of the epitope set. So, we normalized the number of matches by the number of searched epitopes.

For cancer epitopes, the median number of matches per bacterial genus was smaller than that for self epitopes for both Commensals and Neutrals: 0.047 vs 0.053 and 0.062 vs 0.78, respectively (**Figure 5A**), and there was no difference between autoimmune and self epitopes. This trend persisted even when comparing cancer epitopes with original and mutated epitopes, considering only matches with exactly 2 substitutions.

**Figure 5.**
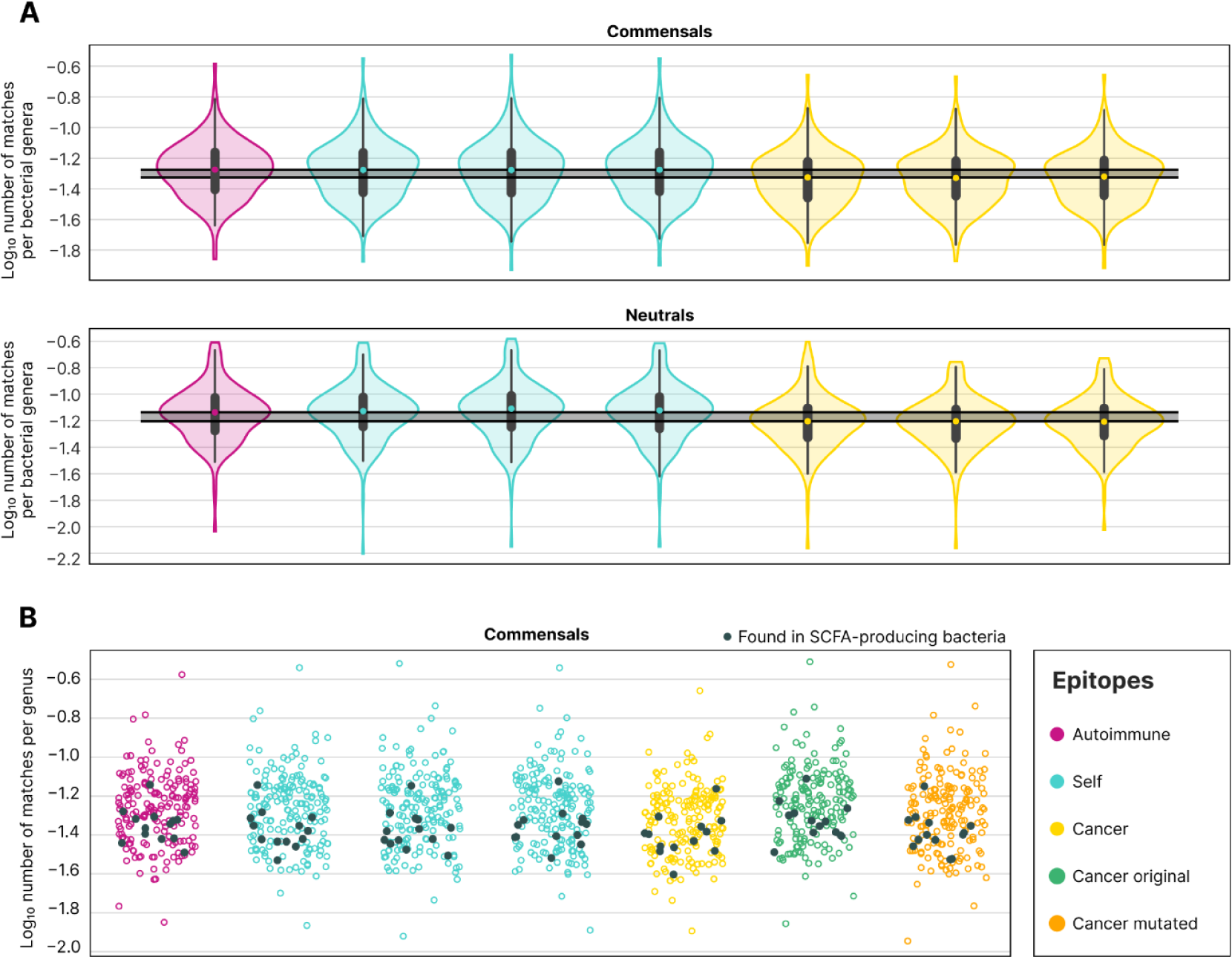
Distribution of the number of matches in proteomes grouped by genus for HLA-I autoimmune, cancer, and self epitopes. **A**: Normalized number of matches found within a bacterial proteome. **B:** Number of matches per proteome for sets of HLA-I epitopes in Commensals and Commensals producing short-chain fatty acids (coloured and black dots respectively). Each dot represents log10 of the number of matches for the proteome of bacterial genus representative normalized by the total number of searched epitopes of this type. The sample sizes are as follows: 25,246, 29,352, and 26,521 epitopes for three sets of normal self; the sample size of the autoimmune epitopes: 9,304 epitopes for the autoimmune set; 4,929, 4,920, and 4,944 epitopes for three sets of cancer neoantigens.

Some commensal bacteria might have a more pronounced tolerogenic effect than others because of the produced metabolites, such as short-chain fatty acids (SCFAs). Thus, we split the studied commensal bacteria into two groups, SCFA-producers and non-producers, and compared the number of matches per bacterial genus in each group. However, the number of matches per genus did not differ between SCFA-producing and non-producing bacteria (**Figure 5B**).

## Discussion

To investigate the frequency of molecular mimicry between T-cell epitopes derived from commensal microbiota and self proteins in this study we estimated the occurrence of different self-epitopes in the proteomes of several groups of microorganisms. Proteomes of commensal bacteria were collected into a group Commensals to measure how close the “extended self” is to our self immunopeptidome. Soil bacteria proteomes were intended to be used for comparison to account for the background proximity of human and bacterial proteomes. The Pathogenes group included proteomes of the pathogenic bacteria, fungi and viruses as a source of antigens presented to immunity in a pro-inflammatory context.

First, we used the immunopeptidome data representing natural HLA-I and HLA-II ligands to estimate which fraction of actual self peptides can be mimicked by peptides derived from proteins of microorganisms. We discovered that a surprisingly high proportion of natural HLA-I ligands (80-90%) and a substantial part of HLA-II ligands (5%) can be detected as potential T-cell epitopes in proteomes of microbiota. As it was previously noted by Trost et al., every human protein contains at least one bacterial pentapeptide or hexapeptide (Trost et al., 2010). Here we focused on those parts of the human proteome that are the source of peptides presented on HLA to be recognized by T cells. For HLA-I, mostly presenting self-antigens, the preferences in the presentation of ligands from specific sets of proteins are well described to date (Karnaukhov et al., 2022; Müller et al., 2017; Pearson et al., n.d.), thus showing that only a small part of human self-peptides are presented to T cells.

To search for the matches for human HLA ligands in bacterial proteomes we used only 9 and 15 aa length epitopes which are the most abundant for HLA-I and HLA-II respectively. Within the search procedure, to account for genetic polymorphisms and the potential TCR ability to recognize similar but not identical epitopes, we allowed up to 2 (for HLA-I) and 4 (for HLA-II) any substitutions (while not affecting HLA-binding) for matches in bacterial proteomes. Both of these restrictions limit our estimation of the similarity between human self and microbial antigen repertoires. Indeed the proportion of the exact matches in our study was substantially lower than matches with substitutions: we detected matches for ∼0.9% of HLA-I and ∼1% of HLA-II ligands. However, while the estimation of the real proportion of such structurally identical epitopes requires development of reliable TCR-pMHC binding predictors, this proportion will be larger than the proportion of exact matches, demonstrating that microbial proteomes can be a substantial source of antigens mimicking human self. Our results complement previous findings that about 0.2% of microbial peptides were identical to human self peptides, and approximately 30% of presented nonself peptides are expected to be indistinguishable from self peptides by T cell receptor when presented in the same HLA context (Calis et al., 2012).

It was shown that the repertoire of self-peptides can be substantially different in normal and inflammatory conditions, as well as between different tissues (Goncalves et al., 2021; Kubiniok et al., 2022). However, we detected no significant prevalence of specific tissues in occurrence of HLA ligands within microbial proteomes. Together with the absence of difference in this respect between Commensals and Neutrals, which are evolutionarily connected to each other, these findings suppose that most of the overlap between human and microbial T-cell epitope repertoire could be related to highly conserved protein sequences, therefore shared between humans and microbes. Nagashima with colleagues demonstrated that T cells mainly target highly conserved in bacteria and abundantly expressed cell-surface antigens of the commensal microbiota. An example of such a conserved antigen in the study was ATP binding cassette transport system, targeted by abundant T cell clonotypes (Nagashima et al., 2023). In our data, we found that 565 bacterial proteins from 62 commensal bacteria containing ATP-binding cassettes have sequences similar to human epitopes.

As molecular mimicry is considered a common cause of post-infectious autoimmunity, this led us to hypothesize that similarity with microbes might elevate the chances of a self epitope to be involved in inflammation. However, commensal microbiota is often subject to peripheral tolerance guided by Treg cells (Cheng et al., 2019) or even central tolerance since some bacterial antigens are presented in the thymus during negative selection (Zegarra-Ruiz et al., 2021). Thus, for a human self epitope, homology with epitopes from commensal or pathogenic (which cause inflammatory response) microbiota might have different consequences.

A direct comparison of the similarity at the T-cell epitope level between humans and commensals and pathogens is unfeasible in our study as the evolutionary relationship within a single sample of microorganisms may have a greater influence on the results than the effect we are investigating. To address this question, we devised an indirect approach. If similarity to commensal microbiota fosters tolerance to an epitope, then epitopes similar to it would have reduced chances to be involved in autoimmunity. On the contrary, epitopes similar to pathogenic microbiota might have elevated chances to be involved in autoimmunity. Cancer uses various methods of immune editing to avoid being noticed by the immune system (Beatty and Gladney, 2015). Therefore, if similarity to pathogens amplifies the risk of detection, cancer might evolve to eliminate such epitopes. If epitopes similar to commensal microbiota promote tolerance, cancer might evolve to acquire them. By comparing the occurrence of predicted human T-cell epitopes from the different sets (autoimmunity-related, cancer neoantigens, and normal self as a control group) in the proteomes of different groups of microorganisms, we are supposed to deduce the effect of similarity with commensals and pathogens.

Contrary to our expectations, we did not observe any differences in results for all microbiota groups (commensals, pathogens, and soil bacteria). For autoimmune HLA-I epitopes, similarity to all of them was slightly lower than for self epitopes. This pattern held even when we isolated and examined short-chain fatty acid-producing bacteria despite their recognized role in promoting tolerance (Kim, 2023). On the other hand, HLA-II autoimmune epitopes displayed greater similarity to each of the three microbiota groups. Results for cancer epitopes also showed the same trend in all microbiota groups: cancer epitopes were less similar to microbiota than self epitopes.

Hence, we have not observed clear and significant differences in the occurrence of different sets of human T-cell epitopes in proteomes of microorganisms which are expected to be differently sensed by the immune system. One of the possible explanations might be the high background level of similarity between human and microbiota T-cell epitope repertoire which is masking the effect. On the other hand, it seems that the potential consequences of similarity at the T-cell epitope level should not be considered in such a mechanistic way. It can not be ignored that the tolerance to commensal antigens should be flexible, allowing the immune system to commensals (Dey and Ray Chaudhuri, 2023; Ost et al., 2021).

We found that cancer epitopes were less similar to microbiota than normal self. To determine if this observation can be attributed to the nature of the original human epitopes mutated during cancerogenesis, we compared similarity levels between cancer epitopes and their original self epitopes without cancer mutation (“cancer original”). They served as a more accurate “self” control. It turned out that the cancer original epitopes were slightly more similar to microbiota than normal self epitopes in our sets, suggestively showing a difference in the number of conserved sequences between cancer original and normal self epitope sets. Indeed, it was shown that during the initial phases, cancer-causing mutations predominantly occur in highly conserved human genes (Piraino and Furney, 2017). This in turn increases the overall similarity level of the entire cancer original set with microbial proteomes due to epitopes derived from such proteins.

To exclude the possibility that the mutations in cancer neoantigens lead to a decreased level of cancer epitope set similarity to microbiota, we randomly mutated cancer original epitopes in concordance with the human mutational profile (Vitkup et al., 2003). Random mutation indeed lowered the previously high similarity of cancer original epitopes to microbiota, but still, it did not reach the levels of cancer epitopes. Thus, it can be explained by the direct evolution of cancer cells to escape similarity with microbiota and thus to escape cross-reactive response from T cells specific to microbial antigens. This is complemented by the recent study that reported sequence homology between some cancer antigens and peptides derived from microbiota, particularly from Firmicutes and Bacteroides (Ragone et al., 2022). It was also shown that molecular mimicry between cancer cells and human microbiota can be deleterious for cancer cells (Fluckiger et al., 2020).

The workflow we used to find differences in the occurrence of epitopes from distinct sets in proteomes of different microorganisms has some substantial limitations. Direct sequence similarity alone does not allow to account for the potential post-translational modifications, the significance of which is increasingly evident in autoimmunity (Anderton, 2004; Zhai et al., 2023). Also, while using HLA-binding prediction we addressed the potential immunogenicity of the epitopes in the sets, the workflow did not consider other factors significantly influencing the immunogenicity of an epitope, such as gene expression level (Chen et al., 2019), cellular localization of a protein (Graham et al., 2018), differences in processing of proteins during antigen presentation (Gomez-Perosanz et al., 2020), protein expression level (Koşaloğlu-Yalçın et al., 2022). However, we obtained similar results between natural HLA-I ligands and predicted epitopes from normal self set, suggesting that all these factors did not significantly affected the results of our study. Also in the absence of reliable and efficient tools for TCR-pMHC binding prediction, we decided to use the very simple and mechanistic approach to account for TCR cross-reactivity by allowing a specific number of mismatches and not considering the physical-chemical properties of amino acid residues. All of these limitations, together with the strong necessity to account for the level of evolutionary conservation between organisms as well as protein sequences (as a source of epitopes), potentially affect the absence of clear differences in the occurrence of distinct epitope sets in the proteomes of different microorganisms and require to be addressed in further research.

To conclude, we observed a high level of similarity between the repertoires of human self and microbial potential T-cell epitopes, which implies that molecular mimicry does not commonly result in autoimmunity. HLA-II autoimmune epitopes showed greater similarity to microbiota than self epitopes, suggesting a potential deleterious effect of molecular mimicry. This is partially supported by the finding that cancer epitopes were less similar to microbiota, suggesting that this might be a result of cancer escape from cross-reactive microbiota-specific T-cells. The results were the same for both pathogenic and commensal microbiota, supposing that tolerogenic or pro-inflammatory effects of molecular mimicry should be investigated with more sensitive approaches.

## Acknowledgments

IVZ was supported by the grant of the Ministry of Science and Higher Education of the Russian Federation grant #75-15-2019-1789.

We thank Dr. Mikhail Shugay and Vadim Karnaukhov for their help with retrieving cancer neoantigens. We are grateful to Dr. Yegor Bazykin for valuable discussions, and Salomé Carcy, Dr. Amitava Banerjee, and Dr. Rishvanth Prabakar for reviewing the draft. We thank Anastasia Troshina for the design of the illustrations.

## Author contribution

VAK – main analysis, contribution to the idea, review and discussion of the results; IVZ – initial idea, study design, review and discussion of the results; VAK - drafting the manuscript; IVZ and VAK – editing and finalization of the text. IVZ – supervising the study.

## Supplementary Figures and Tables

**Supplementary Figure 1.**
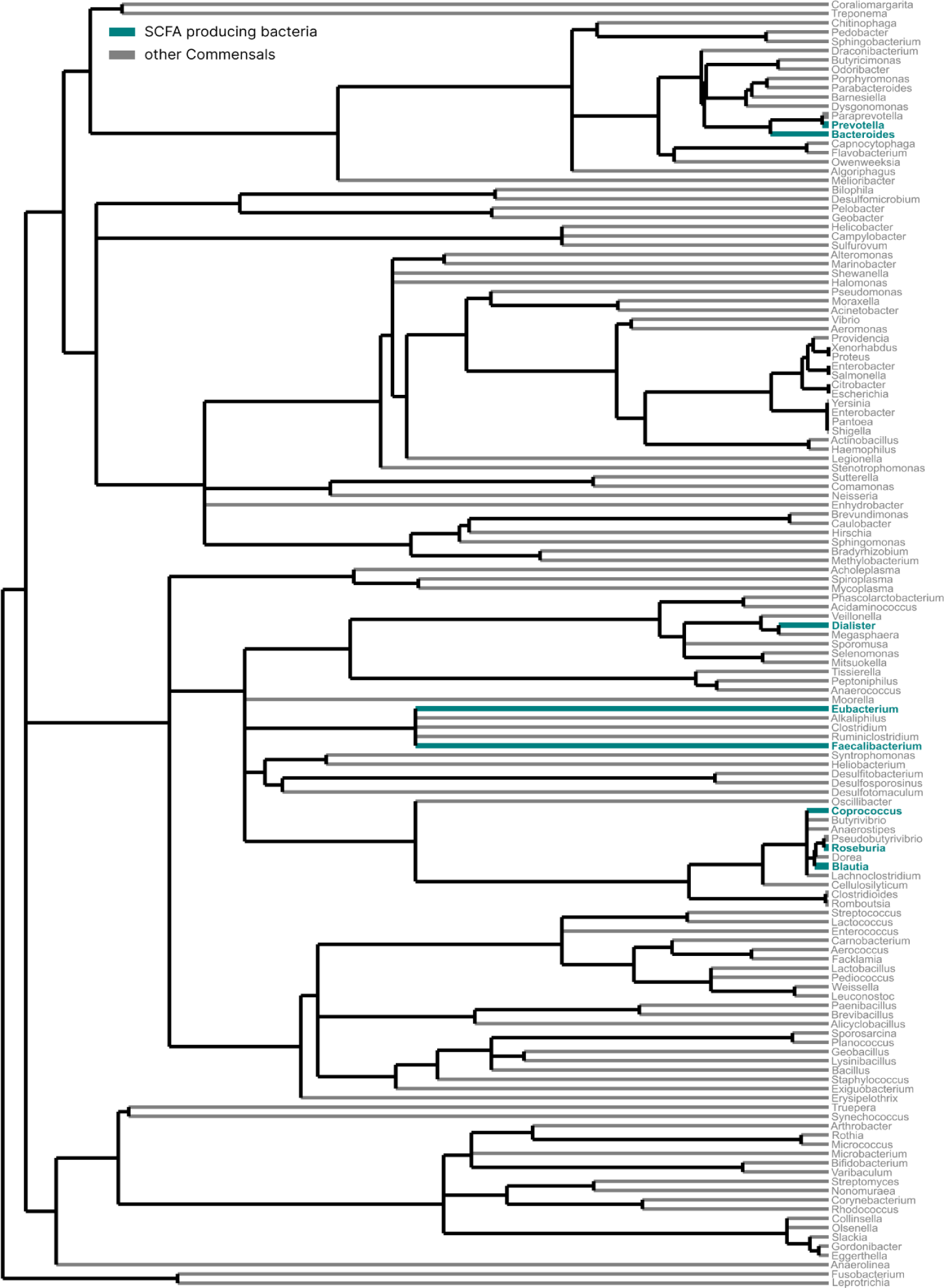
Phylogenetic tree for commensal bacteria. Short-chain fatty acid-producing genera are marked as green. The phylogenetic relationship was obtained from TimeTree (“TimeTree 5: An Expanded Resource for Species Divergence Times | Molecular Biology and Evolution | Oxford Academic,” n.d.).

**Supplementary Figure 2.**
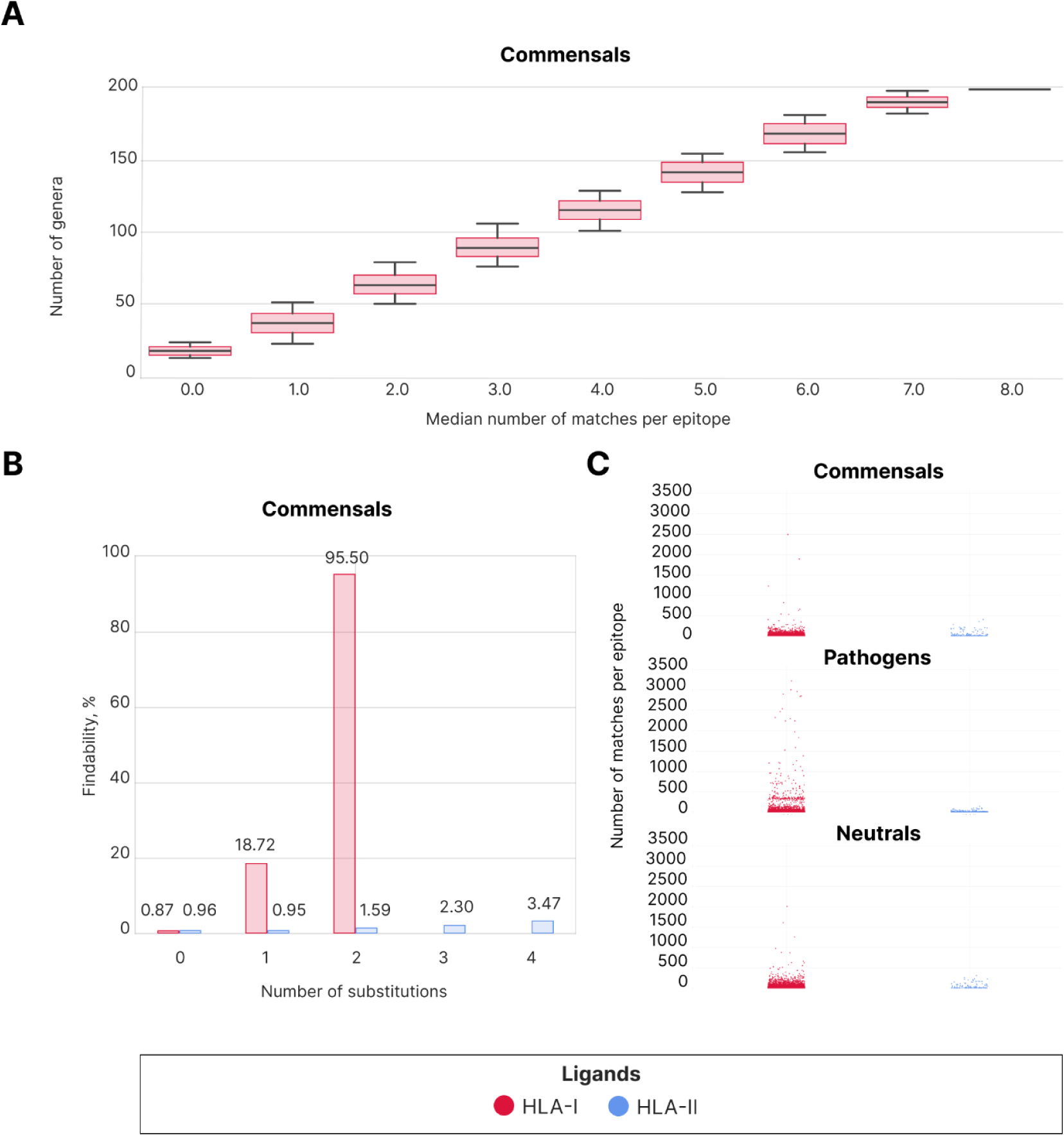
**A**: Median number of matches to HLA-I ligands per genera in random samples from Commensals. **B**: Distribution of findability depending on the number of allowed substitutions in Commensals with the exclusion of HLA-I epitopes which are substrings of HLA-II epitopes. HLA-I ligands are marked as red, and HLA-II ligands are marked as blue. **C**: Number of matches per epitope for HLA-I (red) and HLA-II (blue) ligands in Commensals, Pathogens, and Neutrals.

**Supplementary Figure 3.**
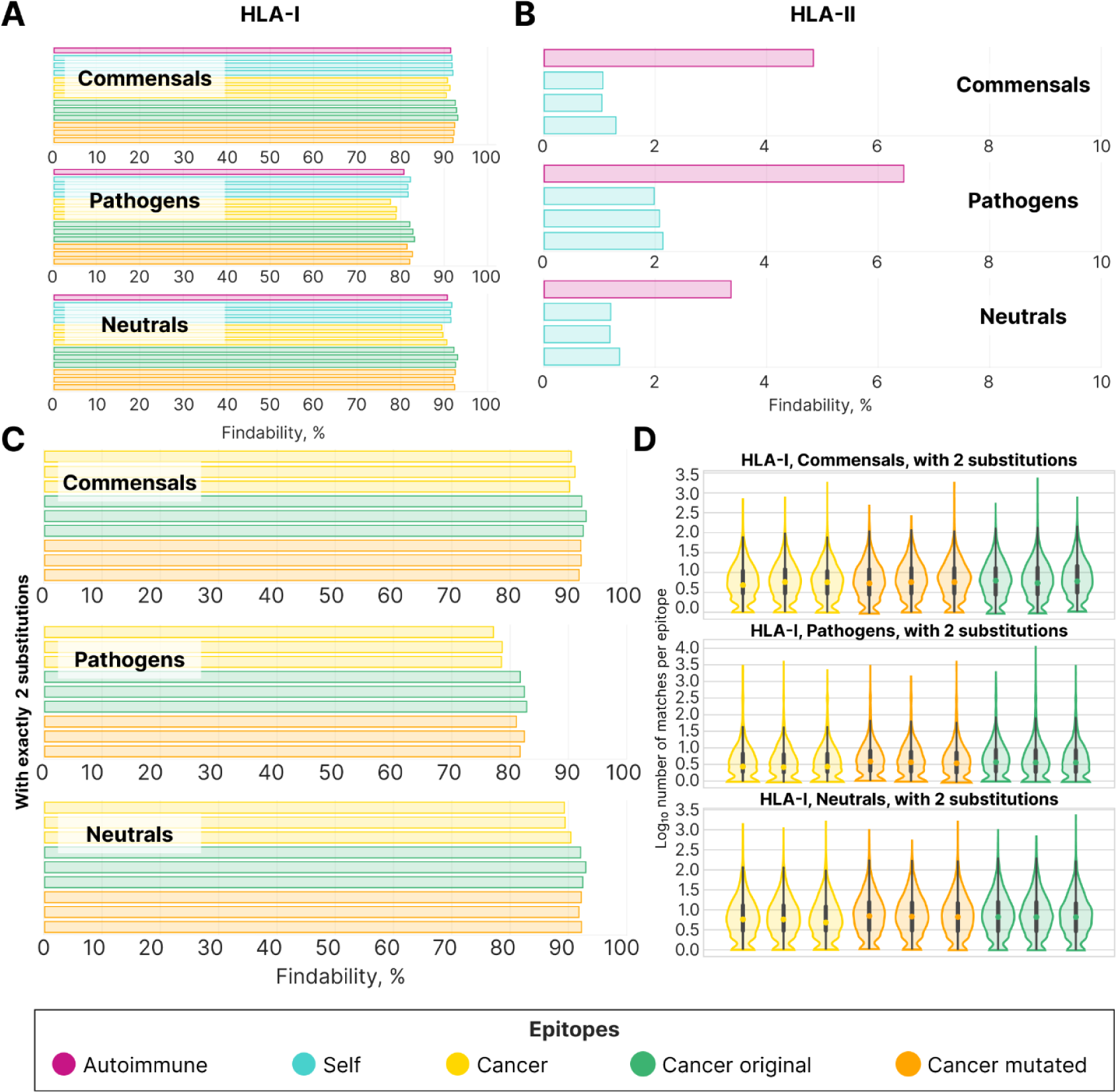
**A**: Findability of HLA-I autoimmune (violet), self (blue), cancer (yellow), cancer original (green), and cancer mutated (orange) epitopes in Commensals, Pathogens, and Neutrals. **B**: Findability of HLA-II autoimmune (violet) and self (blue) epitopes in Commensals, Pathogens, and Neutrals. **C**: Findability of cancer, cancer original, and cancer mutated epitopes in Commensals, Pathogens, and Neutrals. Only matches with exactly 2 substitutions are shown. **D:** Number of matches per epitope for cancer (yellow), cancer original (green), and cancer mutated epitopes (orange) in Commensals, Pathogens, and Neutrals. Only matches with exactly 2 substitutions are shown.

**Supplementary Table 1.** The percentage of random subsamples from self epitopes with a median findability value higher than that for sets of the epitopes of interest (listed in the first column).

|  | Commensals | Pathogens | Soil |
| --- | --- | --- | --- |
| Autoimmune HLA-I | 8.3 | 0.1 | 0.07 |
| Autoimmune HLA-II | 100 | 100 | 100 |
| Cancer -1 | 0.1 | 0 | 0 |
| Original Human -1 | 97.3 | 64.53 | 94.93 |
| Mutated -1 | 93.07 | 22.87 | 85.37 |
| Cancer -2 | 6.6 | 0 | 0 |
| Original Human -2 | 99.53 | 95.07 | 99.97 |
| Mutated -2 | 86.13 | 92.56 | 99.43 |
| Cancer -3 | 0 | 0 | 0.5 |
| Original Human -3 | 99.93 | 99 | 99.6 |
| Mutated -3 | 65.3 | 64.73 | 98.23 |

